# Influence of Mismarking Fiducial Locations on EEG Source Estimation*

**DOI:** 10.1101/544288

**Authors:** Seyed Yahya Shirazi, Helen J. Huang

## Abstract

Mismarking locations of the fiducials can have a significant influence on the digitized electrode locations and cortical source estimation using high-density EEG. Understanding and quantifying how uncertainties in the fiducial locations affect the locations of cortical sources is important for interpreting EEG analyses. We systematically shifted fiducial locations to investigate the relationship between variations of fiducial locations and the corresponding estimations of the source locations. We quantified the uncertainty of the dipole locations using the enclosing volume of the dipole locations and the maximum width of the dipole cluster. Shifting fiducial locations 1.5 cm increased the uncertainty of the dipole locations to span a volume >1 cm^3^ and about 2.5 cm wide. Results suggest that the fiducials need to be digitized accurately within at least 0.5 cm of the absolute actual fiducial location to limit the uncertainty of a dipole location to <1 cm. Additionally, we used random fiducial shift combinations to estimate the effects of combinations of the fiducial shifts on dipole location estimation. This analysis showed that dipole locations were within the bounds of our dipole estimation uncertainty volumes. Based on the outcomes, we suggest marking fiducials carefully before placement of the cap and to use a digitization method with an accuracy of <0.5 cm.

## I. Introduction

Digitizing electrode locations is an essential step for setting up a head model to estimate cortical and subcortical sources from magneto-/electro-encephalography (M/EEG) signals [1]. The digitizing process involves recording three-dimensional positions of the M/EEG electrodes in a global coordinate system and transforming the locations from the global coordinates to the head coordinate system. This transformation requires that the two coordinate systems share at least three anatomical landmarks (i.e. fiducials). The fiducials are typically the left preauricular, right preauricular, and nasion [1, 2].

After digitizing the 3D locations of the fiducials and electrodes, these locations are warped to a head model or vice versa [3]. For studies that involve concurrent tomographic imaging such as magnetic resonance imaging (MRI), digitization, transformation to the head coordinate system, and warping to the head model can be made simultaneously [4], but for other studies, digitized locations must be manually coregistered to construct the head model [1]. Therefore, digitizing can significantly affect the ability to achieve a realistic head model and estimations of source locations [3].

To our knowledge, only a couple of studies examined the effects of digitizing errors on source estimation, and outcomes of these studies do not seem consistent. Beltarchini and collegues [5] suggested that effects of electrode mislocations are negligible on the estimated source locations, while Dalal et al. [6] showed that quality of the output signal degrades significantly with higher uncertainties in the electrode digitization. We did not find any published study that examined the possible effects of mismarking fiducial locations on source estimation.

The purpose of this study was to analyze changes in the estimated source location as a result of shifting the fiducial locations to simulate mismarking of the fiducials. We hypothesized that changes in the locations of the fiducials would have a significant effect on the dipole fitting process and source localization.

## II. Methods

We fitted a mannequin head with a 128-electrode BioSemi (BioSemi B.V., Amsterdam, the Netherlands) cap and digitized the locations of the electrodes and fiducials (left preauricular, LPA; right preauricular, RPA; and nasion, Nz) using an OptiTrack 22-camera motion capture system and a digitizing probe (NaturalPoint Inc., Corvallis, OR). The average (±SD) reliability of this digitization method was <1.50 ± 0.3 mm for digitizing the mannequin head five times.

To define the head coordinate system, we assumed the LPA and RPA were on the X-axis, and Nz was on the Y-axis (Figure 1A). The origin of the head coordinate system was located at the projection of Nz to the X-axis. The Z-axis was defined as the cross product of the X and Y unit vectors and began from the origin. We transformed the digitized electrode locations of the mannequin head to the head coordinate system and used this set of electrodes as the baseline.

**Fig. 1.**
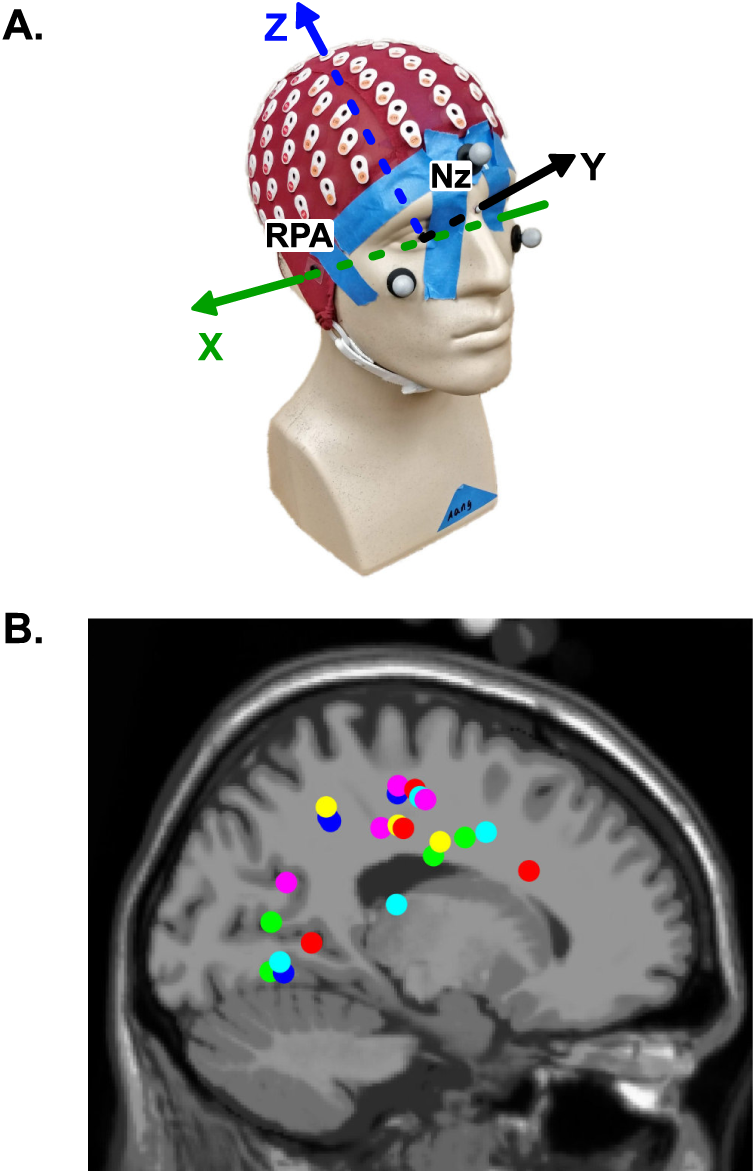
**A.** The mannequin head used for digitization with a representation of the head coordinate system. Right preauricualr (RPA) and nasion (Nz) are labeled in the picture. **B.** The baseline dipole locations of the 23 independent components (ICs) from a separate study but estimated with the mannequin head digitization rather than the individual subject digitizations using ultrasound. These ICs were used to analyze the influence of the systematic fiducial shifts on the dipole locations.

To create multiple sets of electrode locations with various fiducial locations, we shifted one fiducial at a time, in 1 mm increments, up to ±20 mm in the Y and then the Z directions for the preauriculars, RPA and LPA, and in the X and then the Z directions for the nasion, Nz. This process resulted in 12 fiducial shifts per 1 mm increment (3 fiducials × four directions), totaling 240 sets of electrode locations (12 fiducial shifts per 1 mm increment × 20 fiducial shift increments).

Digitization post-processing and the subsequent analyses throughout the study were performed using MATLAB version 9.4 (R2018a, Mathworks Inc., Natick, MA) and EEGLAB [7] version 14.1.2. We used just the fiducial locations to warp the locations of the electrodes and fiducials to the Montreal Neurological Institute’s template head model.

### A. Source Estimation and Dipole Fitting

We used a single representative EEG dataset and weightings of a single Adaptive Mixture Independent Component Analysis (AMICA) from a separate study to perform dipole fitting using the multiple sets of electrode locations with the systematic shifts of the fiducial locations. This representative EEG dataset and ICA had 89 independent components, ICs. We used the DIPFIT toolbox version 2.3 to fit dipoles to the ICs. We visually inspected for dipoles with residual variances <15% that stayed inside the brain area across every fiducial shifts, which resulted in total 23 remaining ICs (Figure 1B). We analyzed the influence of the fiducial shifts on all twenty-three ICs but picked three to illustrate dipole location alterations in detail: 1) the anterior cingulate, 2) the primary somatosensory cortex, and 3) the premotor cortex. Previous studies have shown that these three areas are active in locomotion and error monitoring [8, 9].

### B. Dipole Uncertainty Analysis

We defined the baseline set of dipoles as the dipoles produced using the baseline locations for the fiducials and electrodes (i.e. no shifts). Each set of electrode locations based on a shifted fiducial produced a new set of dipoles. There were 240 sets of electrode locations resulting in 240 sets of dipoles around the baseline dipole. We considered that the spread or size of the cluster of dipoles was representative of the uncertainty related to the resulting dipole locations.

We quantified dipole uncertainty as the volume of a set of tetrahedrons formed from connecting the 12 dipoles created by shifting every fiducial in 1-mm step in every direction (three fiducials × 4 direction/fiducial). We identified the outside boundary of the 12-dipole clusters and created the tetrahedrons using MATLAB’s `convexHull` function.

We also calculated the maximum dipole cluster width between the dipoles with equidistant fiducial shifts (i.e. the same 12 dipoles used for creating tetrahedrons). Since the tetrahedral volumes did not have similar shapes at each fiducial shift, we formed equivalent rectangular cubes with volumes equal to the tetrahedral volumes and the width equal to the maximum cluster width. Hence, for the equivalent rectangular cubes, *V* = *D × E*^2^, where *V* is the tetrahedral volume, which is also the equivalent cube’s volume, *D* is the maximum cluster width that forms the cube’s width, and *E*^2^ is the cross-sectional area of the cube.

We used step-wise polynomial fits to model the relationship between the uncertainty volume and the fiducial shift distance, and the maximum dipole cluster width and the fiducial distance.

### C. Random Fiducial Mismarkings

Creating random combinations of fiducial shifts is another approach to analyze the effects of fiducial mismarkings. We generated 100 electrode location datasets for every 1-mm increment of the random fiducial mismarking combinations, producing 2000 datasets (100 datasets/increment × 20 increments = total 2000 datasets). Each one-hundred mismarkings resided on a circular path with a fixed radius (1 ≤ r ≤ 20 mm) away from the fiducial baseline locations. We ran DIPFIT using each dataset and compared the results with the outcomes of the systematic fiducial shifts.

## III. Results

Shifting LPA, Nz, and RPA in different increments and directions resulted in dipole locations that had curvilinear paths in different planes. Each shift direction created changes in similar directions for the dipoles of the three ICs but with different magnitudes (Figure 2).

**Fig. 2.**
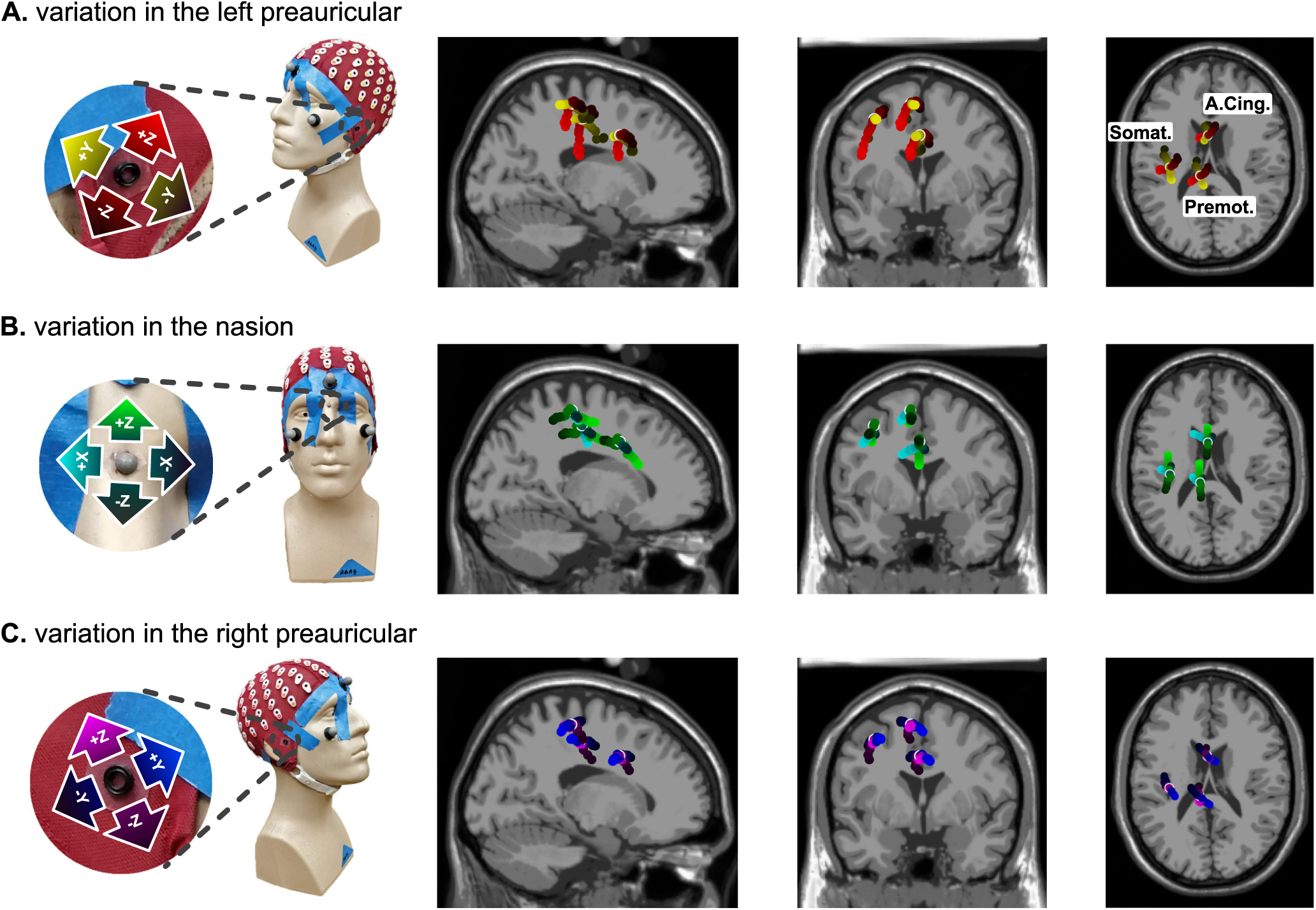
Fiducial mislocations and their corresponding estimated dipoles for **A.** Left preauricular (LPA), **B.** Nasion (Nz) and **C.** Right preauricular (RPA) for three independent components (ICs). Dipoles are located at the anterior cingulate (A.Cing.), the primary somatosensory cortex (Somat.) and at the premotor cortex (Premot.). Lighter colors show positive direction for the fiducial shifts. For each fiducial, resultant dipoles are plotted in sagittal, frontal and top views.

The uncertainty volume increased quadratically as a function of the magnitude of the fiducial shifts (r^2^ = 0.920, Figure 3A). The uncertainty volume was ∼0.06 cm^3^ for fiducial shifts up to 0.5 cm for every IC. For fiducial shifts up to 1.3 cm, all three ICs showed similar increases in uncertainty volumes to 0.5 cm^3^. With fiducial shifts >1.3 cm, the uncertainty volume of all ICs, including the three ICs of interest, began to separate from one another. At a 2 cm shift in fiducials, the average uncertainty volume was >2 cm^3^.

**Fig. 3.**
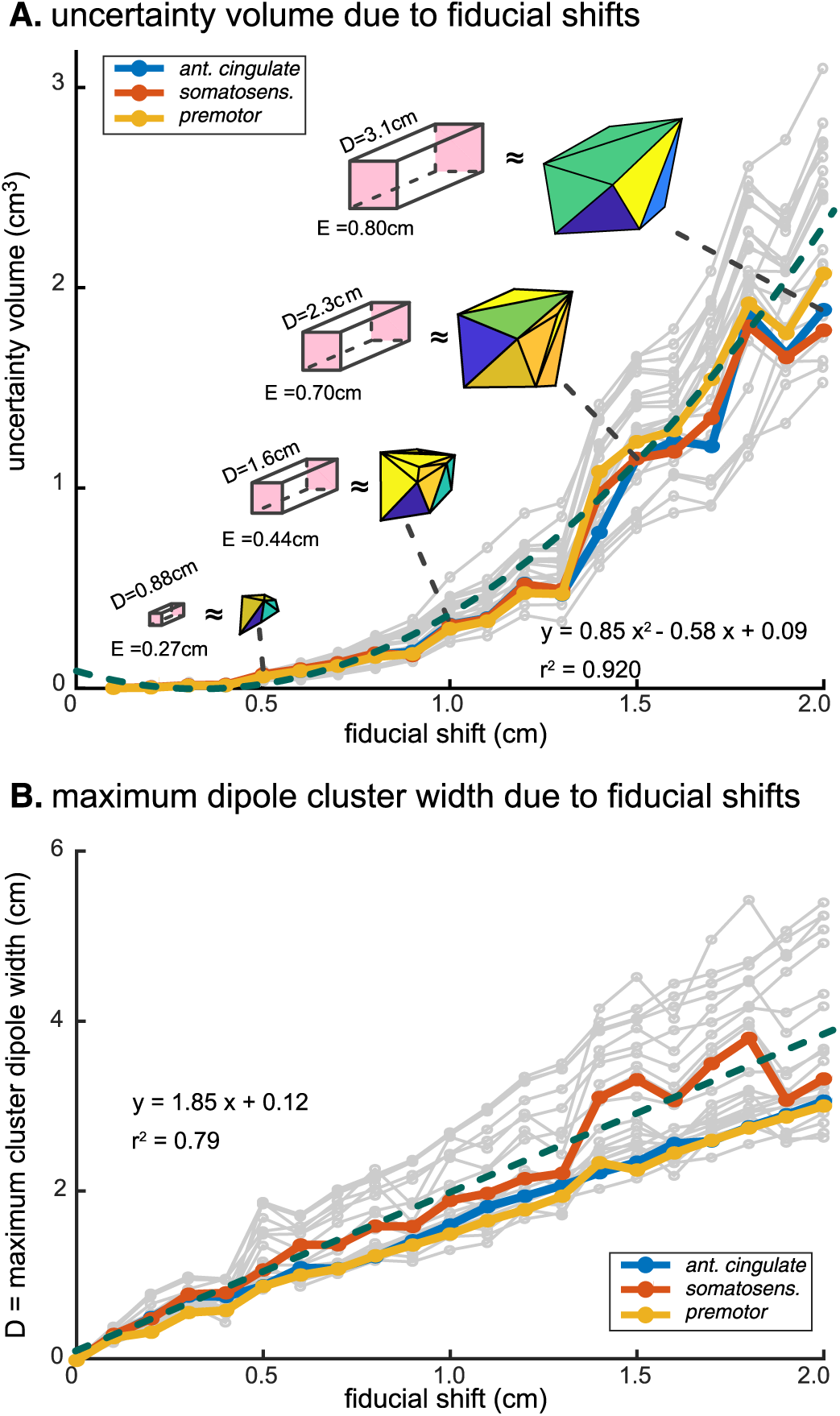
**A.** Relationship between the uncertainty volume and fiducial shifts. ICs in Figure 2 are drawn in color: anterior cingulate (ant. cingulate), somatosensory cortex (somatosens.) and premotor cortex (premotor). Tetra-hedrons represent dipole uncertainty at the anterior cingulate. Pink cubes has the same volume as the tetrahedral volumes, with the same width as the maximum cluster width (D). The green dashed line quadratically models the uncertainty volume as a function of the fiducial shift. **B.** Maximum cluster width (D) of the dipoles estimated from the equidistant fiducial shifts. The green dashed line relates the maximum cluster width to the fiducial shift.

The maximum dipole cluster width for an IC had a linear relationship with the fiducial shifts (r^2^ = 0.79, Figure 3B). The average maximum cluster width exceeded 1 cm for shifts greater than 0.5 cm and was ∼4 cm for fiducial shifts of 2 cm. Some of the cluster widths of different ICs started to deviate from the linear fit for the fiducial shifts >0.5 cm, although, we did not observe similar trend for the uncertainty volumes. The dipole located in the primary somatosensory cortex had larger maximum distances than the ICs in the anterior cingulate and premotor cortex for fiducial shifts greater than 1.3 cm.

Comparing results of the random fiducial mismarking combinations and the tetrahedral volumes from the systematic fiducial shifts, we found that 1) For each 1-mm increment, no random combination of the fiducial shifts (out of 100) could cancel out the effects of the shifts and result in a dipole that could make the uncertainty volume or cluster width smaller, and 2) >95% of the dipole estimations with random fiducial mismarking combinations at each 1-mm increment were in a close proximity of the corresponding tetrahedral surface (within ∼20% of the maximum cluster width).

## IV. Discussion and Conclusion

This study revealed the relationship of fiducial mismarkings during electrode digitization on the subsequent uncertainty of dipole location estimation. We found that shifts of a single fiducial location up to 0.5 cm resulted in an uncertainty volume <0.06 cm^3^ and a maximum distance <1 cm. When fiducial shifts were greater than 1.3 cm, dipole location uncertainty increased to >1 cm^3^ and the maximum distance increased to >2 cm.

One interesting finding was that the largest maximum distances, among the three ICs of interest, occurred in the primary somatosensory cortex, which is an area frequently discussed in the EEG studies related to walking [8, 9]. A previous study found that tangential sources near the boundary of the cortex were more sensitive to electrode location errors, which could explain the larger maximum distances for the dipole at the primary somatosensory cortex compared to a dipole deeper within the cortex such as the anterior cingulate [5].

Another interesting finding was that the linear fiducial shifts mapped to curvilinear dipole paths in different planes, which allowed the use of superposition to estimate dipole uncertainties created from fiducial mismarking combinations. Fiducial mismarking combinations are less likely to cancel each other, since the mismarking directions map to dipole paths in different planes. Fiducial mismarking combinations also created dipole spreads close to the uncertainty volumes that the systematic fiducial shifts had predicted.

Dipole cluster width was more sensitive to the fiducial shifts compared to volumetric uncertainty. Dipole cluster width had a steep linear relation with the fiducial shifts and mismarking the fiducials could change the location of a dipole as much as twice the fiducial mismarking shift. A recent study found that the reliability of a widely-used electromagnetic electrode digitizing system was ∼0.8 cm [10]. Hence, even with perfect markings of the fiducials, there will still be up to 1.5 cm (17% of the head radius) uncertainty for the estimated dipole locations, just as a result of the reliability of the digitizing device.

Limitations of this study were that we did not examine every combination of the fiducial mismarkings and that we only co-registered the fiducials with the MNI head model for warping the electrode locations to the head model. While examining every combination of fiducial mismarkings was not practical, an advantage of analyzing random combinations of the fiducial mismarkings with the same distance from the baseline was that this approach could be used to estimate the mismarking effects for multiple digitizing systems if the digitizing reliability is known. While we could have co-registered more electrodes to the MNI head model, this would lose the individual characteristics of the digitized head. Alternatively, EEGLAB’s NFT toolbox enables warping of the MNI head model to all of the digitized electrode locations [11], but coupling NFT with 240 incremental electrode-location datasets was beyond our computational capacity.

Based on our results, we recommend using a digitizing system with measurement errors less than 0.5 cm and marking the fiducials within 0.5 cm of the actual fiducial to avoid errors greater than 1.5 cm in dipole location. Future work will compare the reliability of different digitizing systems to determine which digitizing systems have measurement errors less than the recommended 0.5 cm. While digitizing the actual locations of the EEG electrodes should provide greater dipole specificity, this study showed that small fiducial mismarkings could result in large dipole location uncertainty. To reduce dipole location uncertainty, care should be taken to minimize the cumulative potential errors from the user mismarking fiducial locations during the digitization process and the measurement errors from the digitization system.

## Notes

* This work was supported by the National Institute on Aging of the National Institutes of Health, under award number 1R01AG054621. The content is solely the responsibility of the authors and does not necessarily represent the official views of the National Institutes of Health.

